# The impact of ants and vertebrate predators on arthropods and plants: a meta-analysis

**DOI:** 10.1101/2022.06.29.498005

**Authors:** Katerina Sam, Marketa Tahadlova, Inga Freiberga, Anna Mrazova, Anna Toszogyova, Rachakonda Sreekar

## Abstract

The trophic interactions between plants, insect herbivores and their predators are complex and prone to trophic cascades. Theory predicts that predators increase plant biomass by feeding on herbivores. However, it remains unclear whether different types of predators regulate herbivores to the same degree, and how intraguild predation impacts these trophic interactions. Specifically, we lack a more comprehensive look at the effects of various groups of predators on a global scale. Here we report a meta-analysis of 486 experiments gathered from 157 publications reporting the effect of insectivorous vertebrates (birds and bats) and ants on abundances of predatory (spiders, ants, others) and herbivorous (chewers and others) arthropods; on arthropod richness and plant damage. Generally, the absence of vertebrate predators led to the increase of predatory arthropods by 18%, herbivorous arthropods by 75%, and plant damage by 47%. In contrast, after the removal of ants, the increase in the abundances of other predatory arthropods did not compensate for missing ants, herbivore arthropods increased their abundances by 53%, and plant damage increased by 146%. The effects of ant exclosures were stronger in communities at lower elevations and latitudes, while we did not detect any clear geographical patterns in the effect of vertebrate exclosures. Neither precipitation nor NDVI had a significant impact on most of the measured effects, and the effect of exclosures was robust for both plant growth forms and different habitat types. We found vertebrate insectivores to be the more dominant predators of arthropods, but we detected that the strength of their trophic cascades was weakened by intraguild predation. On the other hand, we found that although ants were relatively less dominant as predators, and their influence was detectable only in the most productive sites, the effect of trophic cascades on plants they caused was stronger than that of vertebrate insectivores.

## Introduction

Theory on trophic interactions predicts that predators increase plant biomass by feeding on herbivores (Schmitz et al. 2000). Yet, the effect of trophic cascades, i.e., effect of predators on prey and plant communities, has been found to be quite variable (Shurin et al. 2002, Moon and Stiling 2004, Mooney and Linhart 2006), and dependent on ecosystem type (Shurin et al. 2002, Borer et al. 2005, Mooney et al. 2010). Although several meta-analyses have quantified trophic cascades in terrestrial habitats (Schmitz et al. 2000, Halaj and Wise 2001, Mäntylä et al. 2011, Jia et al. 2018), fundamental aspects of how and which predators affect arthropod and plant communities remain unresolved on a global scale.

Interactions between herbivores and predators have been shown to be dependent on changes in the diversity and abundance of communities (Schmitz et al. 2000, Schmitz et al. 2010, Tvardikova and Novotny 2012). These changes may alter the productivity and nutrient flow in the environment which is reflected in trophic cascades (Silliman and Angelini 2012). The changes in species diversity along elevational and latitudinal gradients represent one of the most striking biogeographic patterns on Earth, thus likely affecting the cascades (Rahbek 1995, Colwell et al. 2016). There are several contrasting hypotheses about where we can expect trophic cascades to occur. One of them states that in ecosystems with less diversity and productivity, herbivore populations are more constrained by plant resources than by predatory pressure. In such scenarios, trophic cascades are not likely to occur (Strong 1992). On the other hand, in highly diverse and more productive ecosystems, it should also be difficult to detect trophic cascades, because the effect of predators on plants is diffused in more complex food-webs (Strong 1992). Studies manipulating predator presence across different elevations and latitudes provide an excellent opportunity to determine the effects of trophic cascades on prey populations, and their indirect impact on plants (Schemske et al. 2009).

Vertebrate insectivores such as birds and bats, as well as lizards and frogs in some systems (e.g., Beard et al. 2021), and ants are often considered to be the top predators of arthropods in terrestrial ecosystems. However, it is unclear whether trophic cascades caused by them can affect plant biomass and, if so, in which ecosystems (Schmitz et al. 2000, Mooney et al. 2010). Theory predicts that any direct negative effects of vertebrate insectivores on herbivorous arthropods will be partially or completely offset by the concurrent suppression of arthropod predators, which act as mesopredators of herbivores (Mooney et al. 2010). Nevertheless, extensive empirical evidence exists for the importance of vertebrate insectivores and their top-down effects on trophic cascades (Schmitz et al. 2000, Sekercioglu 2006, Gunnarsson 2007, Whelan et al. 2008).

Other theories suggest that insectivorous arthropods (i.e., ants) do not have strong effects on trophic cascades due to the size-limited consumption of herbivores (Schmitz et al. 2000). On the other hand, bait removal studies suggest that some ant species are able to consume large quantities of arthropods in an ecosystem (Griffiths et al. 2018). Existing predator removal experiments have provided evidence for both the existence and non-existence of trophic cascades between ants and plants (Cerdá and Dejean 2011). Therefore, the ecological question of whether invertebrate insectivores affect trophic cascades between arthropods and plants, and if so, under what conditions, needs to be resolved.

The first publication focusing on the manipulation of predators to evaluate their effect on lower trophic levels was published in 1976 (Solomon et al. 1976), and since then, many similar studies are being conducted every year. Most of these studies have described the effects of vertebrate (birds and/or bats) and ant exclusion separately (Fig. 1). Few exclusion experiments have considered both vertebrates and ants, and even fewer have aimed to exclude lizards (Fig. 1). Recently, factorial experiments treating insectivorous birds and bats as separate variables have started to appear and shed light on their relative effects on arthropods.

**Figure 1.**
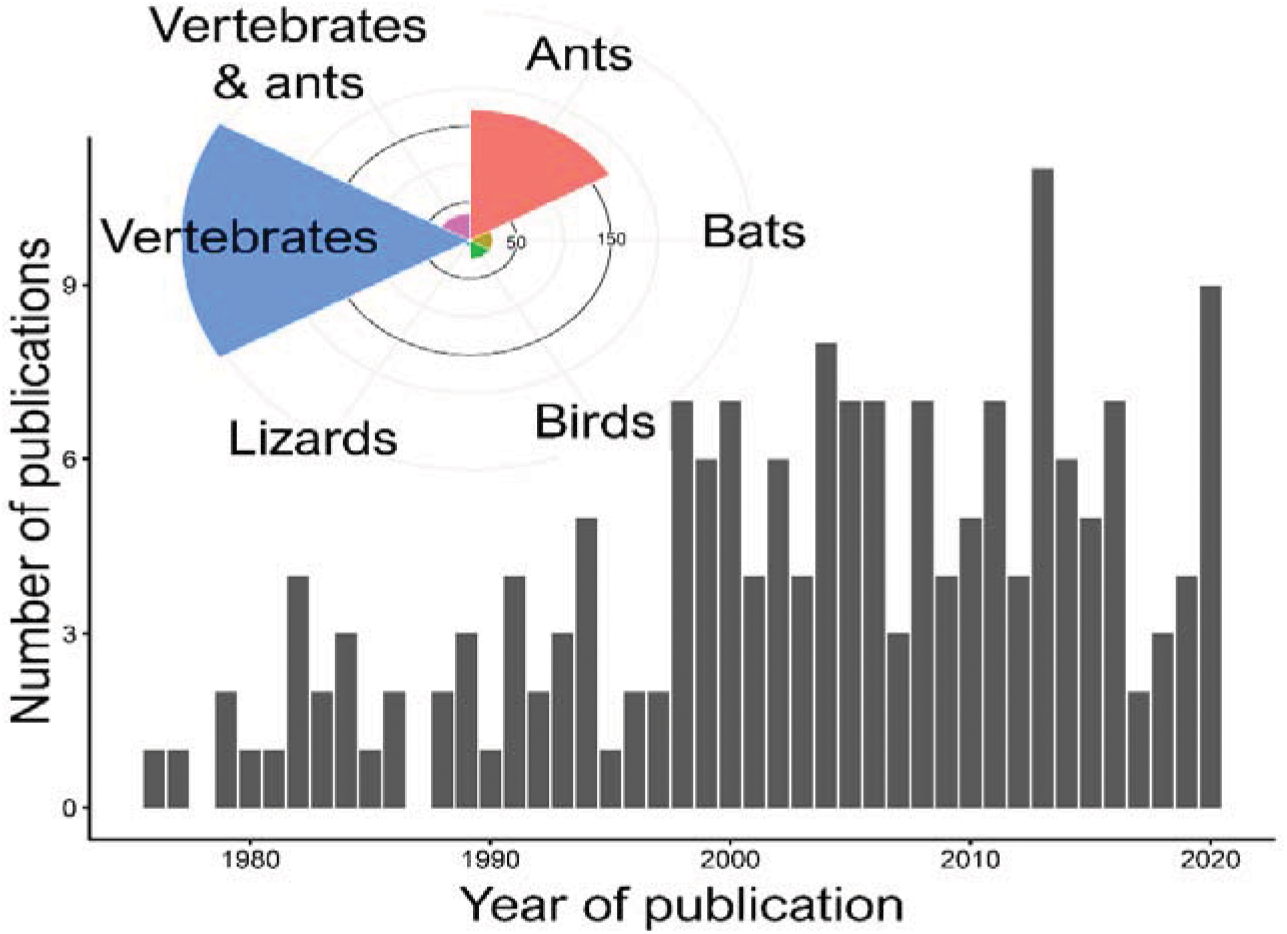
The number of experiments manipulating the presence of predators (vertebrates, vertebrates + ants, ants) published between 1976 and 2019. The bars depict the number of publications manipulating the presence of at least one type of a predator (total = 153). The inset indicates the number of studies excluding individual types of predators. Note that the figure also includes three exclusions of lizards which were not included in later analyses.

To address the knowledge gaps, we conducted a meta-analysis of 486 experiments extracted from 157 publications (including 4 unpublished studies) in which vertebrate insectivores (birds and bats) or ants were excluded. The effects of these exclusions were quantified as changes in the abundance or diversity of arthropods and plant damage. With these data, we tested a set of predictions arising from trophic interaction theories: 1) Generalist vertebrate insectivores should supress both predatory and herbivorous arthropods, and thus, act as intraguild predators in contrast to arthropod predators (i.e., ants) that are more likely to feed exclusively on herbivores; 2) exclusion of vertebrate insectivores should have detectable effects on producers, although these effects might be lower than the direct effect on arthropod communities; 3) exclusion of arthropod insectivores will have no detectable effects on producers, as ants feed mostly on small herbivores that have a low impact on herbivory; 4) we expect predators to have a stronger effect on prey arthropods and plants in more productive environments, at low latitudes and low elevations.

## Methods

We conducted an extensive survey of the literature on manipulative field studies where (i) birds, bats, lizards, and ants were experimentally excluded individually or in combination with either field cages or netting, (ii) matched with appropriate open-access controls, and (iii) the effects of their removal were measured on terrestrial plants and/or naturally occurring arthropod populations or communities. On Web of Science (All collections) and Google Scholar, we searched the terms “bird AND exclosure”, “bat AND exclosure”, “ant AND exclosure” and “lizard AND exclosure”. Our initial search found 546, 18, 19 and 4 studies, respectively. Subsequently, we removed all duplicates and we ended up with 382 potentially suitable studies. We also identified several meta-analyses such as (Mooney et al. 2010) who reviewed 64 studies and (Maas 2019) who reviewed 30 to identify any studies we had missed. We identified a further 153 suitable studies and topped up the dataset with 4 unpublished datasets which we obtained by contacting authors directly (Fig. 2).

**Figure 2.**
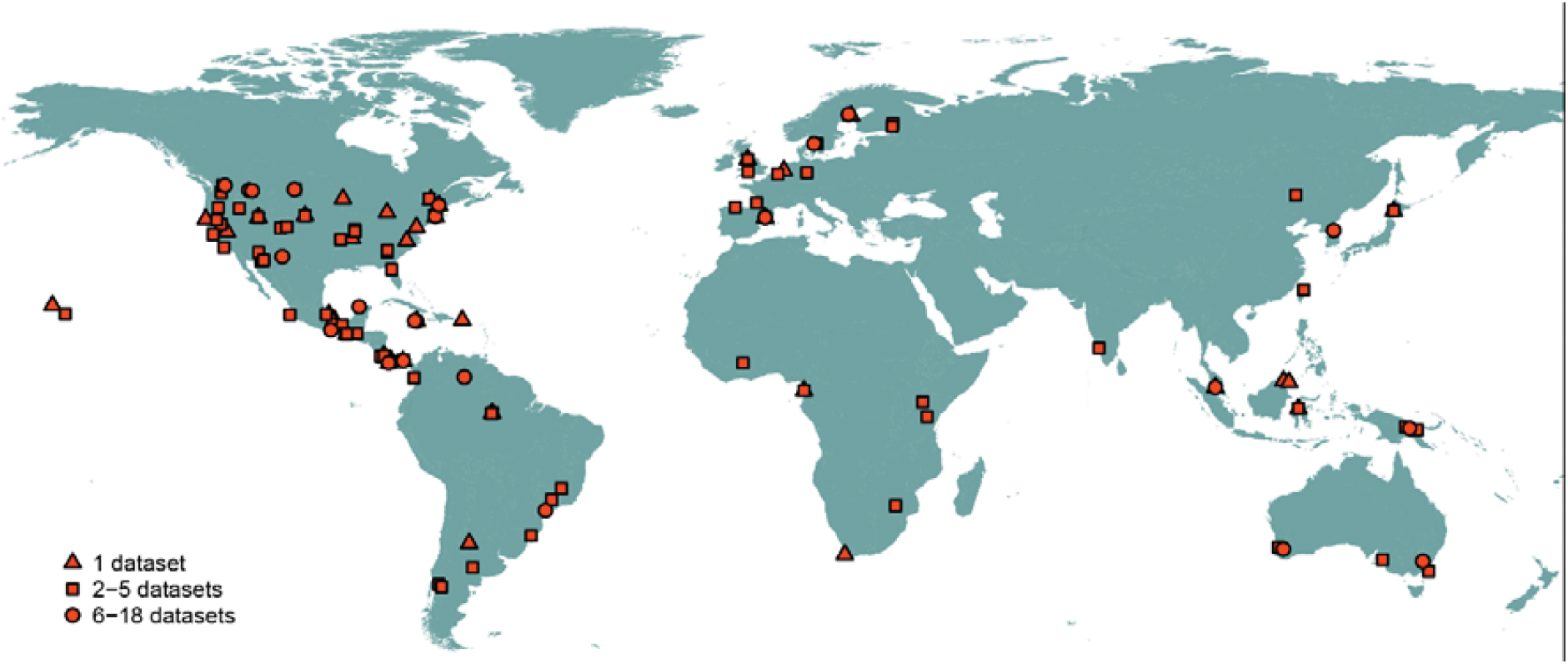
The global distribution of datasets included in the current meta-analysis. Number of datasets (a total of 486) originating from the individual study sites are marked by different shapes. Raw data for individual datasets and sites are available on Github.

For an article to be included, it had to experimentally evaluate the effects of excluding insectivorous predators. Examples of omitted studies were: studies based upon ‘natural experiments’ (or overly correlative studies) (e.g., Jansson and von Brömssen 1981), experimentally staged manipulations that did not use corresponding physical structures excluding predators in the same habitat, such as the introduction of insectivores to islands compared to islands lacking insectivores (e.g., Schoener and Spiller 1999), studies that used acceptable manipulations but measured predation with experimentally deployed sentinel prey rather than natural communities (e.g., Tanhuanpää et al. 2001), studies that only measured effects on pollinators (Muñoz and Arroyo 2004), and studies that measured the effects of birds on aquatic (as opposed to terrestrial) plants (Wootton 1992). Further, we excluded manipulations of predators (lizards N = 9, birds N = 24, and bats N = 24) which did not have enough replicates for robust analyses.

Studies from a single publication were treated as independent when one or more of the following conditions were met: (i) the effects of more than one top predator were measured independently; (ii) the experiments were replicated across multiple field sites; or (iii) when at a single site the effects of a top predator were measured across different ecological contexts, or varying experimental factors (e.g., where birds were excluded on trees with and without ants) or host plant species. In total, we analysed 486 individual experiments from 157 publications in which either vertebrates or ants were manipulatively excluded (data will be available in GitHub and on Dryad upon acceptance).

For each of the study localities, we obtained the Normalized Difference Vegetation Index (NDVI), as an estimate of the environmental productivity based on the spectral properties of vegetation. We used the NDVI data compiled from the MODIS NDVI layers (MOD13A3v006; https://lpdaac.usgs.gov/; Didan 2015) and averaged them for the most productive season of the year (the three most productive months) from the years 2000-2020. The MODIS NDVI data can be acquired at https://lpdaac.usgs.gov/products/mod13a3v006/. The data we obtained had a spatial resolution of 1 km, which represented a suitable area range around the data points of the studies.

### Data analysis

The top-down effects of ant and vertebrate exclusions: We determined the effect of exclusion type (two explanatory variables – ant or vertebrate insectivore exclosure) by changes in the abundance of all arthropods, herbivorous arthropods, predatory arthropods, arthropod richness, and plant damage measured as chewing herbivory on foliage (i.e., response variables) using a linear mixed-effect model. Later, we investigated the effect of exclusion at a finer scale (i.e., finer response variables) by calculating changes in the abundances of chewing herbivores, ants, spiders, and other predatory arthropods. Finally, we determined the effect of exclusion by calculating changes in the ratios between (i) all predatory arthropods and all herbivorous arthropods, (ii) all predatory arthropods excluding ants and all herbivorous arthropods, and (iii) all predatory arthropods excluding ants and chewing arthropods. Tests for all hypotheses were based on unweighted natural log response ratios LRR (Curtis and Wang 1998, Hedges et al. 1999) calculated from mean responses in the presence or absence of a predator (ant or vertebrate). LRR is an effect size measurement that quantifies the results of experiments (i.e., response variables) as the log-proportional change in between the means of the variable in treatment (in the absence of insectivorous predators, □_I-_) and in control group (in the presence of insectivorous predators, □_I+_) and was thus calculated as (ln[□_I-_/ □_I+_]). To compare the ratio of mesopredators (i.e., predatory arthropods) to herbivores in the presence (P:H_I+_) and absence (P:H_I-_) of insectivorous predators, we used a log response ratio as ln([P_I-_:H_I-_/ P_I+_:H_I+_). It is important to note, that we calculated LRR the classical way (as ln(experimental mean/control mean); e.g., Curtis and Wang 1998, Hedges et al. 1999), and our results thus differ from similar reviews. Specifically, from those where reverse LRR was used as ln(control mean/experimental mean (Mooney et al. 2010), or where log response ratio was calculated as lnR = ln(control mean) − ln(experimental mean) (Mäntylä et al. 2011).

In our meta-analysis, the results of the multiple treatments were sometimes extracted from a single “study”. Therefore, we accounted for the effect of a “study” by specifying it as a random variable in our models. The studies used in in our analyses sampled our response variables for varying durations (0 to 90 months). We therefore also accounted for the effect of “duration” by specifying it as a covariate in the model. We built two global models to determine effects of exclusion on the response variables:

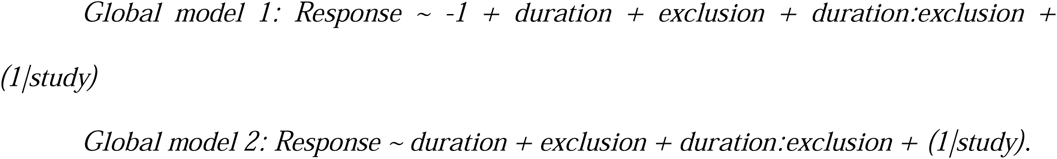

The first global model considers the true intercept to be 0 and the second global model does not. Therefore, the first global model compares each exclusion type with the control, whereas the second global model compares the two exclusion types against each other. We resampled 10000 times from the posterior estimates of the model to compute their 95% confidence intervals, and p-values were computed using the method described in. We used a backward elimination technique to select the best model (i.e., minimal adequate model) to determine the effects of exclusion on the response variables.

To calculate the effects of environmental variables on exclusion treatments we used individual models to determine if treatment effects varied across environmental gradients for the measured response variables. We examined the effect of elevation, latitude, precipitation, habitat type and plant growth form on treatment effects. These variables were extracted from the publications where possible or from WorldClim databased based on the provided GPS location.

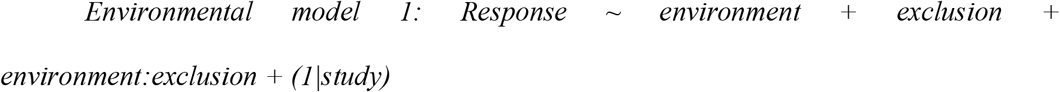

Following our earlier results, we included duration in models where response variables were affected by study duration.

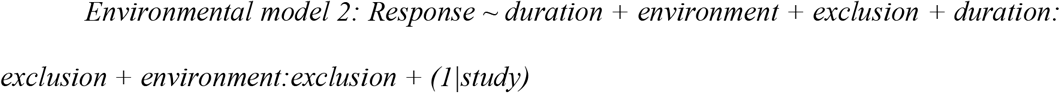

Furthermore, for models that examined the effects of elevation on the response variable, we used latitude as the first variable to account for latitudinal differences across studies. Similarly, for models that examined the effects of latitude on the response variable, we used elevation as the first variable to account for elevational differences across studies.

### Summary of the dataset

The datasets we analysed spanned a latitude of -41.2 to 64.23 degrees (mean ± SE = 19.82 ± 3.32) and altitudes of 0 to 3700 m a.s.l. (650.4 ± 150). The experiments were more often conducted in natural than in agricultural landscapes (342 vs. 144, respectively). The majority of the studies were conducted on mature vegetation (238 experiments) in contrast to saplings (93 experiments) or mixed vegetation (11 experiments). Experiments conducted on trees (308) were more common than experiments on shrubs (178). Studies from tropical and temperate regions were represented almost equally (188 vs. 221 experiments) while 61 experiments were conducted in subtropical regions and only 16 in boreal regions. The shortest experiments lasted 7 days and the longest lasted 96 months (9.56 ± 1.1 months). Enclosed areas ranged from 0.1 m^2^ to 1000 m^2^ (35.14 ± 1 m^2^).

We identified that the description of the treatment was a major weakness in many of the studies. In 255 experiments, the authors reported that they excluded birds. However, in all of them, the netting was placed permanently around the plant, so the treatment actually excluded both birds and bats, and potentially also lizards. The effect of birds and bats were only studied separately in 39 experiments, in which nets were removed at either daytime or night-time. Surprisingly, only 151 of the analysed experiments were finalized by complete sampling of all arthropods. In contrast, arthropods were only partially surveyed in 204 experiments (usually only for chewing herbivorous insect). In 162 experiments, there was no account of arthropods and only herbivory was surveyed. Further, there were only 125 experiments that provided detailed information on which orders of arthropods were affected. From these studies, we extracted information about the impact of exclusion treatments on spiders (101 experiments), ants (60 experiments), chewing arthropods including Coleoptera, Orthoptera and the larvae of Lepidoptera and sawflies (Symphyta, Hymenoptera; 125 experiments).

## Results

According to our expectations, the total number of arthropods increased significantly after the exclusion of insectivorous vertebrates (hereafter vertebrates; (P < 0.001, Fig. 3a, Table S1). In contrast, ant exclosure did not lead to significant changes in the total number of arthropods (P = 0.084; Fig. 3a, Table S2). The contrast between the two treatments was significant (P = 0.045). The absence of vertebrates and ants led to an increase in arthropods by circa 69% [natural log ratio LRR = 0.53 ± 0.23 (mean ± 95%CI)] and by 12% (LRR = 0.11 ± 0.49), respectively, although the effects of ant exclusion were insignificant (Table S2). The trends in the results remain the same even after we controlled for the effect of ant exclosure and worked only with a subset of studies that provided data on individual orders of arthropods. In ant exclosure treatments, the total abundances of arthropods other than ants did not change significantly (P = 0.642). We also did not find any significant effect of experimental duration on the difference in total abundances of all arthropods in both treatment types (Figure S1, Table S1).

**Figure 3.**
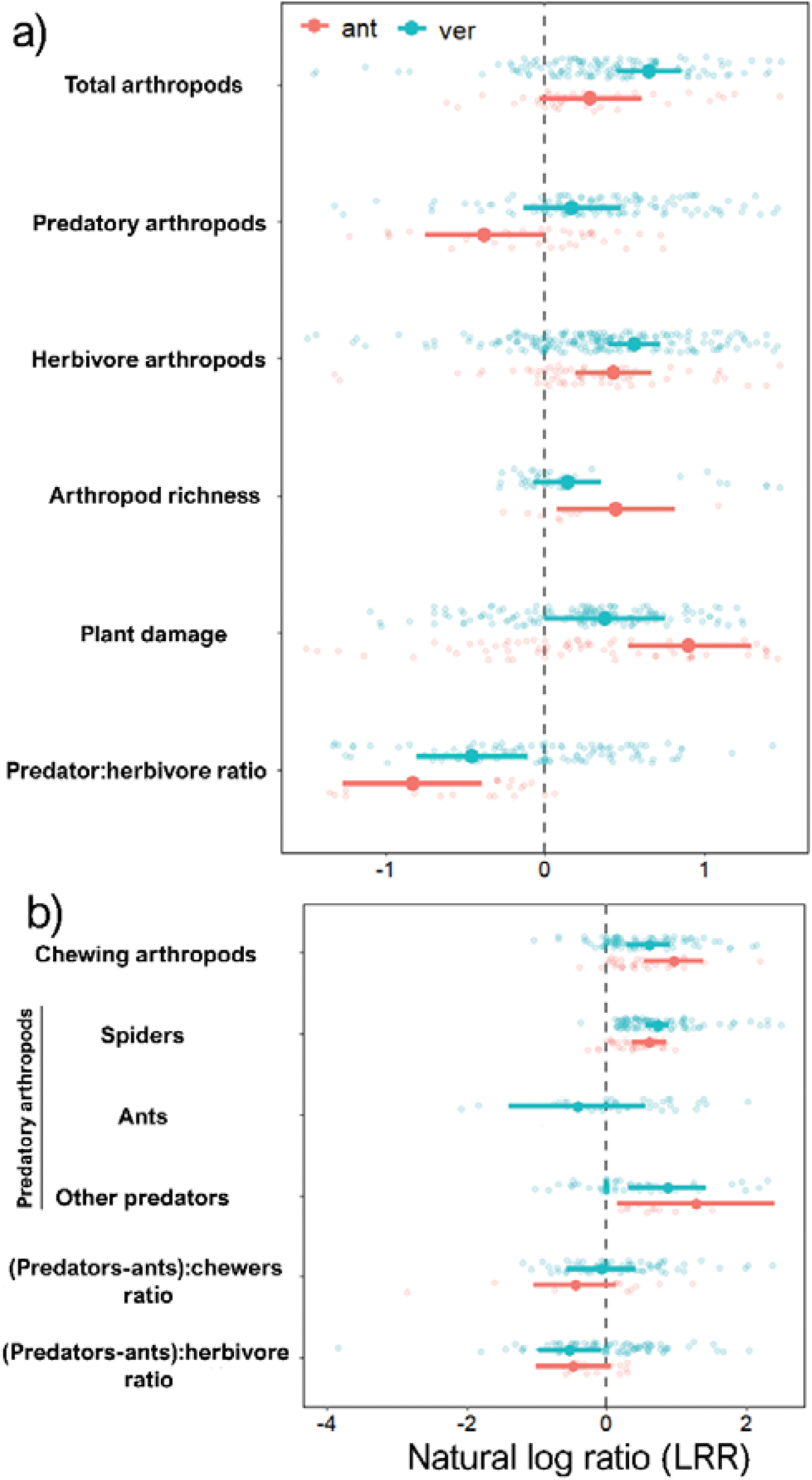
Caterpillar plot showing how the total abundance of all arthropods, predatory arthropods, herbivore arthropods, arthropod richness, plant damage and predator-herbivore ratio respond to ant (red) and vertebrate (green) exclusion treatments (a). Based on the subset of studies providing more detailed data, it also shows how chewing arthropods, spiders, ants, other predatory arthropods, and ratio between predators without ants and chewing arthropods and the ratio between predatory arthropods without ants and herbivore arthropods respond to ant (red) and vertebrate (green) exclusion treatments (b). Following our previous results, we included duration in individual models where response variables were affected by study duration (refer to Figure S1, Table S1). The X-axis shows the mean effect sizes of natural log ratios (LRR = ln(exclosure/control)) with 95% CI of individual response variables. Effect size values above zero indicated that the absence of predators due to the exclusion treatments is harmful to plants, as the abundances of arthropods in exclosures increased and potentially caused increased herbivory damage to plants. Note the different scales on the x-axes in part (a) and (b).

Arthropod species richness was not significantly affected by the type of exclosure (P = 0.085, Table S1). The resulting arthropod species richness increased significantly when ants were excluded (by 57.4%; P = 0.015) but did not change when vertebrates were excluded (by 15.6%; P = 0.201; Figure 3a, Table S2). The duration of the experiment again had no effect on changes in species richness (Figure S1, Table S1).

More specifically, the treatments had significantly different effects on all predatory arthropods (P < 0.001, Fig. 3a, Table S1). The abundances of predatory arthropods increased non-significantly by 18% in vertebrate exclosures (LRR = 0.17 ± 0.30; P = 0.264) in comparison with the respective control treatments (Fig. 3a, Table S2). In contrast, their abundances tended to decrease in ant exclosures by 31.2% (P = 0.055; LRR = -0.37 ± 0.36, Fig. 3a, Table S2). This was mainly result of the successful removal of ants as the part of the treatment (by 98%; LRR = -4.23 ± 2.86; P < 0.001, Table S1, Table S2) and by the fact, that such loss of ants in ant exclosures was not fully compensated by the increase in the abundances of spiders (by 84%, LRR = 0.61 ± 0.25, Table S2) or other predatory arthropods (by 263%; LRR = 1.29 ± 1.12, Table S2). Despite their abundances increasing significantly in comparison with control treatments (Table S1), the natural abundances of other predators were typically much lower than the abundances of ants and spiders. Surprisingly, we also detected a decrease in the abundances of ants in vertebrate exclosures (by 33%, LRR = -0.41 ± 0.56, P = 0.401, Table S2). The increase of abundances of individual groups of predatory arthropods did not differ significantly between vertebrate and ant exclosures (Table S1). Specifically, the abundances of spiders increased significantly by 107% (P < 0.001; LRR = 0.72 ± 0.16) and the abundances of other predatory arthropods increased significantly by 137% (P = 0.002; LRR = 0.86 ± 0.56) (Table S2) in vertebrate exclosures.

For the abundances of herbivorous arthropods, there was no significant difference between the two treatment types (P = 0.239, Fig. 3a, Table S1). Abundances of herbivorous arthropods increased significantly (P < 0.001) both in vertebrate and ant exclosures by 75% and 53%, respectively (Fig. 3a, Table S2). Similarly, as in the case of the abundances of all arthropods, we did not find any significant effect of the duration of the experiment on the abundances of predatory and herbivorous arthropods (Figure S1, Table S1). Even looking at chewing herbivores separately did not reveal any significant change between the two treatments (P = 0.194), despite the exclusion of vertebrates leading to an increase in chewers by 161% and exclusion of ants leading to an increase by only 84% (both these increases were significant, P < 0.001) (Table S2).

The change in ratio between all predatory and all herbivore arthropods was affected by the duration of the experiment (P = 0.004) in addition to the type of predator excluded (P = 0.036) (Table S1). After accounting for the variation in study duration by including it as a covariate in the model, the difference between the two treatment types was still significant (P = 0.029). Both the exclusion of ants (P < 0.001; decrease of the ratio by 56%) and the exclusion of vertebrates led to an overall significant decrease (P = 0.012, decrease of the ratio by 36%) in the ratio between predatory arthropods and herbivorous arthropods (Fig. 3a). However, as we showed above, this might be purely a methodological aspect. After limiting the data to ant exclosure studies which provided detailed arthropod data, the effect between predatory arthropods without ants and all herbivores did not change significantly (P = 0.082; LRR = -0.48 ± 0.59, decrease of the ratio by 38%) and the ratio between predatory arthropods without ants and chewers decreased non-significantly by 36% (P = 0.134; LRR = -0.45 ± 0.59, Fig. 3b, Table S2).

Similarly, the ratios remained unchanged in the vertebrate exclosures (Table S1). Specifically, after the exclusion of vertebrates, the ratio between predatory arthropods without ants and all herbivores did not change significantly (P = 0.182; decrease of the ratio by 41%), similarly to the ratio between predatory arthropods without ants and chewing herbivores (P = 0.149; decrease of the ratio by 7%) (Table S2). These observed decreases in the ratios in exclosures were typical in situations where abundances of herbivores increased, while abundances of predatory arthropods decreased only slightly in exclosures in comparison to controls.

Plant damage was significantly affected by the duration of the experiments (P = 0.008, Figure 3a), and the influence of duration on plant damage varied significantly between treatment types (P = 0.012; Figure S1; Table S1). The duration of the experiment had a weak significant effect on the vertebrate exclusion treatment (P = 0.041) but interacted strongly with the ant exclusion treatment (P < 0.001). Plant damage reduced with the duration of experiments and dropped from a ca. 113% increase in herbivory (LRR = 0.759) in studies excluding ants for less than 3 months, to an increase by 19% (LRR = 0.18) in studies excluding ants for longer than 11 months.

After accounting for variation in duration by including it as a covariate in the model, there was no significant difference between the two treatment types (P = 0.712). Both vertebrate exclusion (P = 0.051) and ant exclusion (P = 0.001) led to significantly higher herbivorous damage than on control plants (by 45% and 146% respectively; Fig. 3a, Table S2). We did not conduct any robust analysis on the response of plant biomass, and we found no correlation between herbivorous damage and plant biomass response in the experiments where both biomass and herbivorous damage were measured (N = 34, r = 0.19, P = 0.29).

Latitude had no significant effect on any, except two, of the measured response variables (Fig. S2, Table S3). There was a significant interaction between latitude and treatment in both arthropod species richness and abundances of herbivore arthropods (Fig. S2, Table S3). After the removal of ants, arthropod species richness increased more at low than at high latitudes and no trend was detected in the exclosures of vertebrates (Fig. S2, Table S3). Abundances of herbivores tended to increase more at high latitudes than at low latitudes in ant exclosures, while the opposite pattern was generally found in vertebrate exclosures (Fig. S2, Table S3).

Along the elevational gradient, several significant patterns were detected. Increases in the abundances of herbivore arthropods, = chewing arthropods, species richness of arthropods and plant damage were significantly higher at lower elevations than at higher elevations in ants exclosures, while there was no pattern, or only a slight pattern detected in vertebrates exclosures (Fig. 4B-D, L; Table S3). The abundances of all predatory arthropods, spiders, and the ratio of predators without ants to herbivore arthropods increased more at low than at high elevations, and this pattern was detected in both treatments (Fig. 4E, F, J; Table S3).

**Figure 4.**
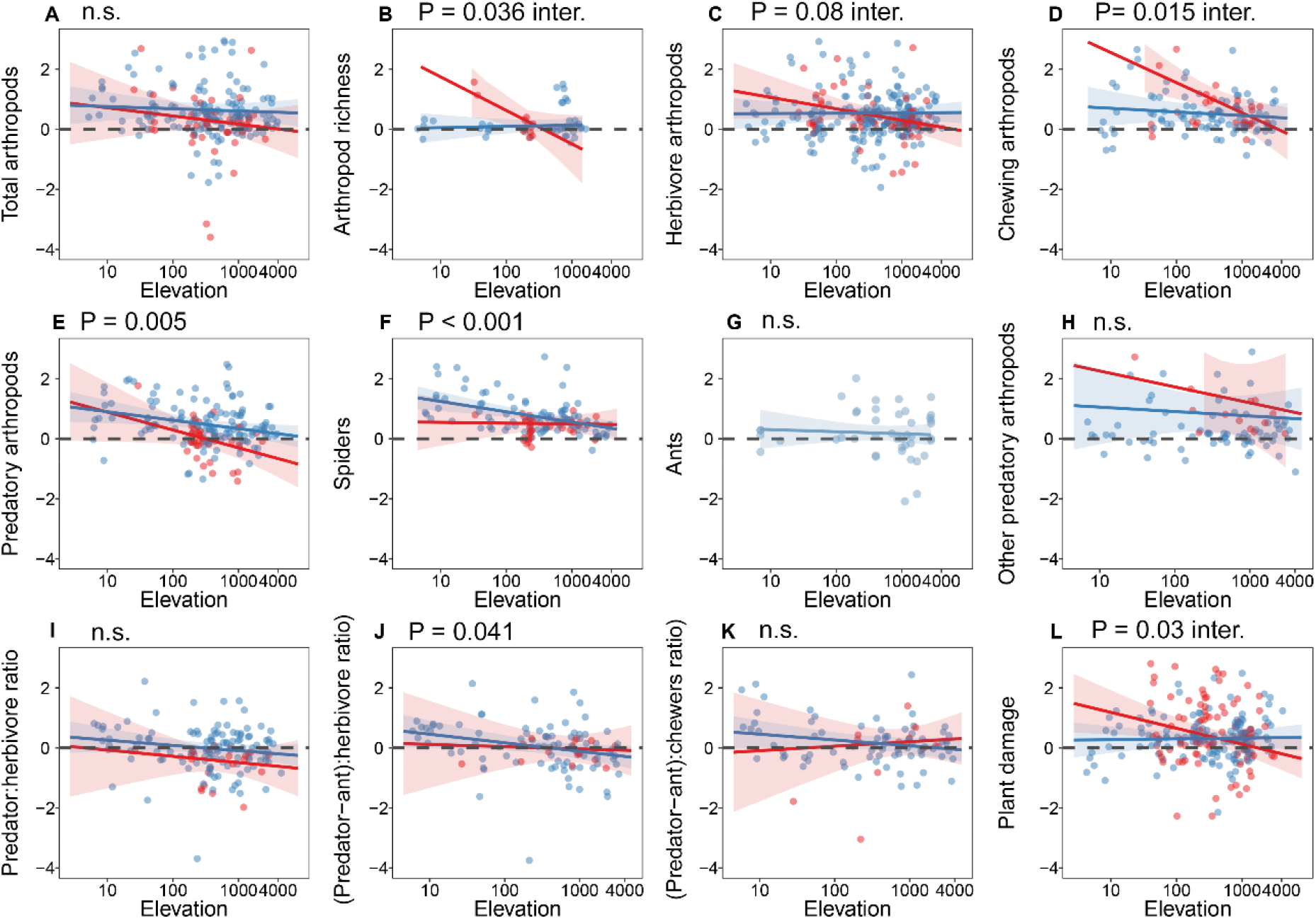
The effect of elevation and treatment (exclusion of ants in red, exclusion of vertebrate insectivores in blue) on response variables. Elevation had no significant effect on abundances of all arthropods (A), ants in vertebrate exclosures (G), other predatory arthropods (H), the ratio between all predatory arthropods and all herbivore arthropods (I) and on the ratio between predatory arthropods without ants and herbivore arthropods (K). Elevation had a significant effect on the abundances of all predatory arthropods (E), spiders (F) and the ratio between predatory arthropods without ants and herbivores (J). The interaction between elevation and treatment (ant or vertebrate exclosures) had a significant effect on arthropod species richness (B), abundance of herbivore arthropods (C), chewing arthropods (D) and plant damage (L). Y-axes shows the natural log ratios (LRR = ln(exclosure/control)) of individual response variables. Log-scaled elevation was used in models and the actual elevation was plotted on the x-axes to ease the visual interpretation of the results. Latitude was used in all models as the first covariate. The complete results of all models are in Table S2.

Precipitation did not have any effect on many of the measured response variables. However, there was an increase in the richness of arthropods with increasing precipitation after ants were excluded (Fig. 5B). Further, the increase in the abundance of herbivore arthropods was higher in drier than wetter habitats when ants were excluded, but the opposite trend was observed in vertebrate exclosures (Fig. 5C). The abundances of spiders increased with increasing precipitation both in vertebrate and ant exclosures (Fig. 5F). The results for precipitation were actually very similar to those using NDVI (Table S4, Fig. S3) as an explanatory variable and the subsequent analysis revealed significant (P < 0.001) correlations between precipitation and mean NDVI for productive seasons (r = 0.73) and between precipitation and the median NDVI of monthly values (r = 0.65). However, when considering either of the two NDVI measurements, the abundances of spiders increased only slightly and non-significantly with increasing productivity (Table S4, Fig. S3). When considering the median monthly values of NDVI across the whole year, the increase of plant damage in ant exclosures was significantly stronger in more productive sites than in less productive sites (Table S4, Fig. S3).

**Figure 5.**
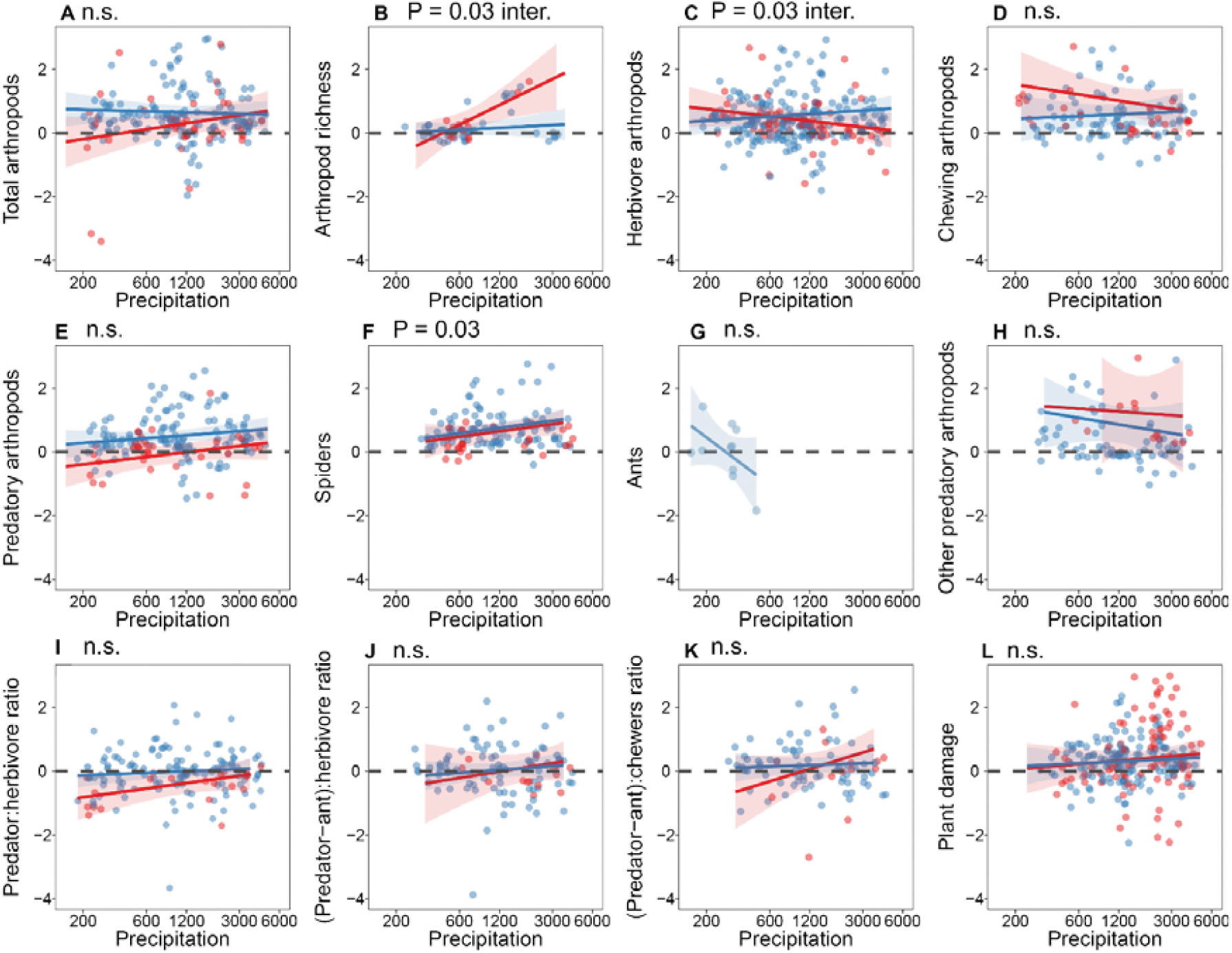
The effect of precipitation and treatment (exclusion of ants in red, exclusion of vertebrate insectivores in blue) on response variables. Precipitation had a significant effect on arthropod richness and abundances of herbivore arthropods in interaction with treatment (B, C). Abundances of spiders increased more in places with higher precipitation than in places with lower precipitation (F). Y-axes show the natural log ratios (LRR = ln(exclosure/control)) of individual response variables. Log-scaled precipitation was used in models and actual precipitation was plotted on the x-axes to ease the visual interpretation of the results. Latitude was used in all models as the first covariate. The complete results of all models are in Table S4.

The effects of exclosure treatments generally did not change between natural and agricultural habitats (Fig. S4, Table S5). However, the abundances of herbivore arthropods increased significantly more in natural habitats than in agricultural habitats (Fig. S4). A similar, yet non-significant, trend was seen in chewing arthropods (Fig. S4). Further, there was an interaction between treatment and habitat type, suggesting that vertebrate exclosure led to a lower increase of spiders in ant exclosures than in vertebrate exclosures in agricultural habitats, but that pattern was reversed in natural habitats (Fig. S4).

The observed results were also robust for plant growth forms (Fig.S5, Table S5). Yet, species richness increased significantly more on saplings than on mature trees in both treatments. The abundances of chewing arthropods, predatory arthropods and the ratio between predatory arthropods and chewers changed in interaction with growth forms (Fig.S5, Table S5). Overall, the responses to vertebrate exclusion tended to be stronger on mature trees than on saplings, while the effect of ant exclosure was often higher on saplings than on mature trees. Also, the abundances of ants in vertebrate exclosures increased on mature trees but decreased on saplings (Fig. S5, Table S5).

## Discussion

Consistent with our first hypothesis, vertebrate insectivores supressed the total abundances of arthropods, and both predatory and herbivore arthropods individually. Yet, potentially imbalanced feeding by vertebrate insectivores resulted in an unexpected difference between the abundance of predatory and herbivorous arthropods, as their abundances increased by 18% and 75%, respectively, in the absence of vertebrates. It is important to note that even balanced feeding, during which the predatory and herbivorous arthropods are consumed equally, would preclude further predation from them on herbivores, thus leading to some, likely low, increase of herbivorous arthropods. We believe that this mechanism can contribute to the observed results, but it might not be the main driver. Regardless the mechanisms, our results differ from previous meta-analyses, which have reported a more balanced effect of insectivorous predators on predatory and herbivorous arthropods, resulting into increase of their abundances by 38% and 39% respectively in vertebrate exclosures (Mooney et al. 2010). However, our results are consistent with several case studies that reported increases in the abundance of herbivorous arthropods by ca. 70% and mesopredators by ca. 20% in the tropics (e.g., Floyd 1996, Bridgeland et al. 2010, Schwenk et al. 2010, Tahadlova et al., in prep.).

In contrast to vertebrate insectivores, ants supressed herbivorous arthropods slightly more than predatory arthropods. Dietary studies further demonstrate that vertebrate insectivores consume both predatory and herbivorous arthropods in nearly equal proportions, whereas ants feed predominantly on phytophagous arthropods (Spiller and Schoener 1990, Halaj et al. 1997, Gunnarsson 2007, Iakovlev et al. 2017, Roslin et al. 2017). This would imply that only vertebrate insectivores, but not ants, act as intraguild predators. Yet our data did not support these results strongly and these preliminary results set the stage for our subsequent hypotheses.

Contrary to our expectation that the effects of ants on plants might be undetectable because they are likely to feed on smaller arthropods, vertebrate exclusion actually caused weaker cascades to plants than ant exclusion, highlighting the importance of mesopredators in ecosystems. Since ants seemed to consume mainly herbivorous arthropods, the exclusion of ants resulted in an increase in herbivory damage by ca. 147%. In contrast, despite having strong negative effects on the abundance of predatory arthropods, insectivorous vertebrates also suppressed herbivorous arthropods and exerted an overall beneficial effect on plants, reducing plant damage by ca. 47%. Our result for insectivorous predators is only slightly higher than a previous estimate showing that vertebrate insectivores reduced plant damage by 40% globally (Mooney et al. 2010). Our findings thus seem to be in direct contrast to empirical and theoretical studies suggesting that smaller predators (i.e., ants) generate weak or no trophic cascades, and that the loss of large predators (i.e., vertebrates) has greater consequences than the loss of smaller predators in otherwise similar ecosystems (Laws and Joern 2013, DeLong et al. 2015). While both types of exclosures have similar effect on herbivore abundances, ant exclosure have greater negative effect on plants. However, another mechanism might be in play. First, most of the ant exclosure studies were conducted in systems where ants have a tight mutualistic relationship with plants. Thus, in these systems, excluding ants might naturally have a stronger positive cascading effect on herbivores and herbivory which is not reflected in other ecosystems. Second, the observed positive effect might also be influenced by ants having strong, mutualistic associations with hemipterans, which we were not able to address in this study. A lack of ants might lead to a decline in aphid communities, positively affecting chewing herbivores (Mooney et al. 2006).

We expected that damage caused by herbivores would accumulate over time (Ritchie and Penner 2020) and that we would observe a stronger effect on plant herbivory than on arthropod abundance (Sobek et al. 2009). Unexpectedly, we observed a rather similar effect of predator removal on both herbivory and arthropod abundances. These results may suggest a progressive weakening of top-down effects (Schmitz 2008), which is supported by our observation that plant damage decreased over time for the duration of the experiment, going from an increase of 113% in 3 months to an increase of only 19% in 11 months. Analysing experiments of various lengths might have resulted in the dilution of a potentially existing effect. We suggest that the length of experiments be well designed and discussed in future experiments and analyses.

Effect of ant exclusion was affected by precipitation, or by NDVI, which was very closely correlated with precipitation, whereas the effect of bird exclusion did not change across the precipitation gradient. However, in contrast to our fourth expectations, the effect of vertebrate predator exclusion did not change with latitude (Roslin et al. 2017). Similar discrepancies in the predation by arthropod and vertebrate insectivores along a latitudinal gradient were found by a global study using dummy caterpillars as a proxy for predation rates (Roslin et al. 2017). In this study, we show that this trend, or lack thereof, is detectable at lower trophic levels in plant herbivory. The observed results are expected, as vertebrate insectivores are nearly ubiquitous, and their overall species richness (Kissling et al. 2012) and abundance (Emlen et al. 1986) does not seem to decrease dramatically with latitude. Further, earlier studies comparing temperate and tropical locations showed that bird predation is similar between sites (Remmel et al. 2011), and that ant predation is significantly higher in the tropics than in temperate areas (Jeanne 1979, Novotny et al. 2006).

Along the elevational gradient, predator exclusion has the most significant effect on predatory arthropods. After the exclusion of both vertebrate insectivores and ants, the abundance of predatory arthropods increased by ca. 250% in lowlands (up to 200 m a.s.l.), whereas no increase was detected in uplands. We observed a similar trend in herbivorous arthropods after ants were excluded, but not when vertebrate insectivores were excluded, likely leading to a deviation from normal ratios of predatory to herbivorous arthropods. Predatory arthropods are usually spiders and hymenopterans, which have naturally higher abundances and richness in lowlands than in highlands (Lalisan et al. 2015, Ribeiro et al. 2019). It is therefore expectable that they were able to better fill the empty niches after the top predators disappeared from the lowlands (Halaj et al. 1997). Further, arthropod diversity increased in the lowlands but not in the uplands after ants were excluded. This pattern might be a statistical artefact, where diversity of arthropods increases after high abundances of a social insect species are removed from a sample by the treatment. Alternatively, ants might function as a strong competitive agent and their removal actually led to an increase in diversity as new insect species occupied new niches. Changes in plant damage after the removal of ants, as discussed above, were more detectable in lowlands than in uplands. Specifically, herbivory damage increased by ca. 600% in lowlands but only by ca. 100% in uplands after ants were excluded.

We also detected an interaction between habitat type and predator type that suggests a stronger effect of insectivorous vertebrates in natural habitats, while the effect of ants was stronger in agricultural habitats. In natural habitats, the absence of insectivorous vertebrates led to higher (by ca. 290%) abundances of arthropods than the absence of ants (by ca. 150%). In agricultural habitats, the abundances of arthropods increased by ca. 250% and ca. 690% respectively (insectivorous vertebrates vs. ants). Despite the large changes in the abundances of arthropods, the increase in plant herbivory in the absence of predators was detectable only in natural habitats. In previous studies, ants exhibited top-down effects by limiting arthropods and increasing plant growth in agricultural landscapes (Schmitz et al. 2000, Philpott and Armbrecht 2006). Our study found the same trend but did not confirm that ants could significantly reduce plant damage in agricultural habitats. This is likely because our results are not robust enough (16 of the 35 ant exclosure studies in agricultural localities did not measure herbivory damage), and they were again affected by the duration of the experiments (long studies reported a decrease in herbivory by about 110% and shorter studies an increase of ca. 30%). We showed that ants have a stronger effect on herbivore abundances on trees than they do on shrubs. In theory, trees could represent a higher-quality microhabitat because of higher productivity in canopies than in shrubs and understories. However, the currently available data might not be suitable for the resolution of this hypothesis, as only a few of the experiment analyses here were conducted in the canopies of natural forests, and the majority of the canopy studies were conducted in agricultural lands such as coffee or cocoa plantations. Even when studies were conducted in undisturbed forests, the experiments were done in the lower stories of the crowns as there was limited access to upper strata of the forests. Future studies should therefore focus on the importance of predators for forest productivity.

Summarizing our results, we show that vertebrate insectivores supress both predatory and herbivorous arthropods to such an extent that the trophic cascades to plants are hardly detectable. The observed, little to none, effect exhibited by the exclusion of vertebrate insectivores is surprising in the view of current theories (Vance-Chalcraft et al. 2007) but our results show they are not extensible to predict the effects of vertebrate insectivores on these terrestrial systems. On the other hand, the effect of ants on herbivorous arthropods, cascaded to plants relatively strongly. Our analyses further detected predicted associations between the effect of ant removal and precipitation (or NDVI) but not latitude. We failed to detect these predicted positive associations for vertebrate insectivores. Although we have documented some clear patterns in top-down control and the cascading effects of ants and vertebrate insectivores, the limitations of existing data prevent a definitive conclusion. We argue that more studies measuring the effects of predators on all arthropod feeding guilds, and on herbivory damage are needed to ascertain stronger conclusions. Future studies should also be conducted over larger scales than most of the existing studies and over longer periods of the time.

