## Supplement for "The impact of ants and vertebrate predators on arthropods and plants: a meta-analysis"

Supplementary material: The impact of ants and vertebrates on arthropods and plants: a meta-analysis

**Figure S1.** Effect of duration of the experiment and treatment (exclusion of ants in red, exclusion of vertebrate insectivores in blue) on response variables. Duration of the experiment did not have significant effect on the effect of exclosure on total abundance of arthropods (A), arthropod richness (B), abundance of herbivore arthropods (C), chewing arthropods (D), all predatory arthropods (E), abundances of spiders (F), abundances of ants after the vertebrate insectivores were excluded (G), and on abundance of predatory arthropods other than spiders and ants (H). Duration of the experiment had significant effect on ratio between all predatory arthropods and all herbivore arthropods (I), and on ratio between predators without ants and all herbivores (J). Duration itself and in interactions with the treatment (ant or vertebrate exclosure) affected ration between predators without and chewing arthropods (K) and plant damage (L). Y-axes show natural log ratios (LRR = ln(exclosure/control)) of individual response variables. Note different span of y-axes. X-axes show duration of the experiments, spanning from 0.25 of month to 96 months. Log-scaled duration was used in models and actual duration of the experiment in months was plotted on log-scale to ease the visual interpretations of the results. Complete results of all models are in Table S1.

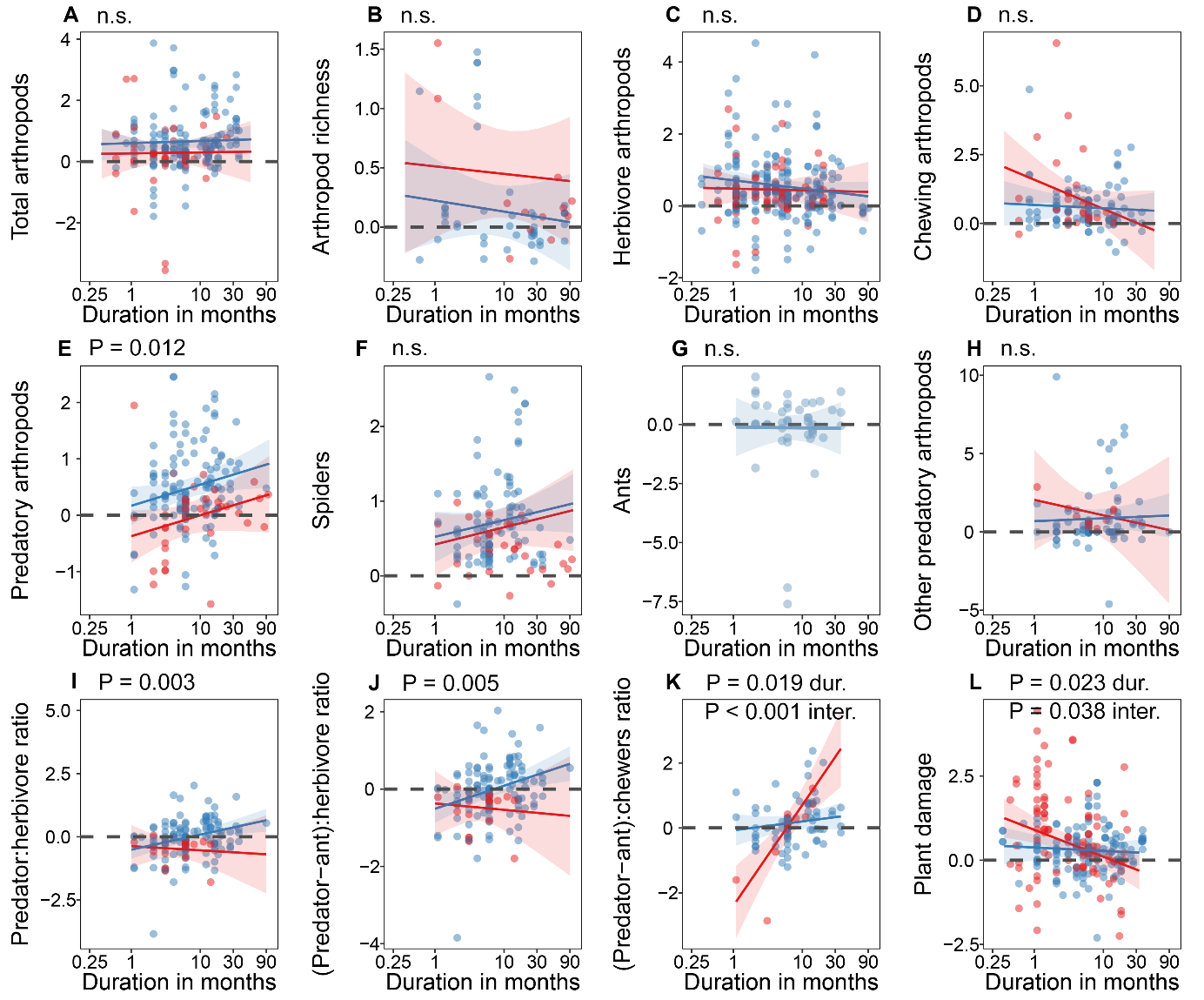

**Table S1.** Results of the models considering the effect of duration and the type of the treatment (ant or vertebrate exclosure) on response variables. The effects of the duration are visually shown in Fig. S1.

|  |  | Chisq | Df | P |
| --- | --- | --- | --- | --- |
| Total arthropods | Duration | 0.1067 | 1 | 0.744 |
|  | Treatment | 4.0249 | 1 | **0.045** |
|  | Duration * treatment | 0.0071 | 1 | 0.933 |
| Arthropod diversity | Duration | 0.4067 | 1 | 0.524 |
|  | Treatment | 2.9609 | 1 | 0.085 |
|  | Duration * treatment | 0.0117 | 1 | 0.914 |
| Herbivore arthropods | Duration | 1.9856 | 1 | 0.159 |
|  | Treatment | 1.3891 | 1 | 0.239 |
|  | Duration * treatment | 0.3979 | 1 | 0.528 |
| Chewing arthropods | Duration | 1.0636 | 1 | 0.302 |
|  | Treatment | 1.6886 | 1 | 0.194 |
|  | Duration * treatment | 1.9734 | 1 | 0.160 |
| Predatory arthropods | Duration | 6.2578 | 1 | **0.012** |
|  | Treatment | 15.687 | 1 | **<0.001** |
|  | Duration * treatment | 0.0001 | 1 | 0.993 |
| Spiders | Duration | 2.8436 | 1 | 0.092 |
|  | Treatment | 0.6293 | 1 | 0.428 |
|  | Duration * treatment | 0.0005 | 1 | 0.983 |
| Ants | Duration | 0.0127 | 1 | 0.910 |
|  | Treatment | 30.1031 | 1 | **<0.001** |
|  | Duration * treatment | 0.0727 | 1 | 0.787 |
| Other predators | Duration | 0.0186 | 1 | 0.891 |
|  | Treatment | 0.5448 | 1 | 0.460 |
|  | Duration * treatment | 0.3352 | 1 | 0.563 |
| Predatore:herbivore ratio | Duration | 8.325 | 1 | **0.004** |
|  | Treatment | 4.3929 | 1 | **0.036** |
|  | Duration * treatment | 1.5826 | 1 | 0.208 |
| (Predators-ants):herbivore ratio | Duration | 7.7346 | 1 | **0.005** |
|  | Treatment | 0.0426 | 1 | 0.836 |
|  | Duration * treatment | 0.1447 | 1 | 0.704 |
| (Predators-ants):chewers ratio | Duration | 5.4434 | 1 | **0.019** |
|  | Treatment | 0.1023 | 1 | 0.749 |
|  | Duration * treatment | 14.6245 | 1 | **<0.001** |
| Plant damage | Duration | 6.9629 | 1 | **0.008** |
|  | Treatment | 0.6002 | 1 | 0.438 |
|  | Duration * treatment | 6.3752 | 1 | **0.012** |

**Table S2.** Natural log ratios (LRR, mean and low and high confidence intervals) for all response variables and treatments as predicted by the minimal adequate models. Percentual change is then calculated as inversed function from LRR (EXP(log)*100-100) and the results of the simulations (as P values) testing whether the change in the two treatments differed significantly from 0 (b).

| Response | Excluded | Change by % | Predicted LRR | Low CI | High CI | Change of LRR from 0 (P value) |
| --- | --- | --- | --- | --- | --- | --- |
| Total arthropods | ant | 11.9 | 0.113 | -0.373 | 0.603 | 0.084 |
|  | ver | 68.6 | 0.522 | 0.283 | 0.758 | **<0.001** |
| Arthropod richness | ant | 57.4 | 0.454 | 0.088 | 0.817 | **0.015** |
|  | ver | 15.6 | 0.145 | -0.071 | 0.361 | 0.201 |
| Predatory arthropods | ant | -31.2 | -0.373 | -0.744 | -0.006 | 0.055 |
|  | ver | 18.3 | 0.168 | -0.132 | 0.469 | 0.264 |
| Ants | ant | -98.6 | -4.239 | -5.642 | -2.869 | **<0.001** |
|  | ver | -33.6 | -0.410 | -1.397 | 0.560 | 0.401 |
| Spiders | ant | 84.0 | 0.610 | 0.360 | 0.854 | **<0.001** |
|  | ver | 106.8 | 0.727 | 0.565 | 0.890 | **<0.001** |
| Other predators | ant | 262.0 | 1.286 | 0.161 | 2.404 | 0.272 |
|  | ver | 137.7 | 0.866 | 0.310 | 1.428 | **0.002** |
| Herbivore arthropods | ant | 53.2 | 0.426 | 0.190 | 0.667 | **<0.001** |
|  | ver | 75.2 | 0.561 | 0.397 | 0.722 | **<0.001** |
| Chewing arthropods | ant | 160.9 | 0.959 | 0.541 | 1.381 | **<0.001** |
|  | ver | 83.9 | 0.609 | 0.303 | 0.909 | **<0.001** |
| Predator:herbivore ratio | ant | -56.3 | -0.827 | -1.263 | -0.397 | **<0.001** |
|  | ver | -36.4 | -0.453 | -0.808 | -0.108 | **0.012** |
| (Predators-ants):herbivore ratio | ant | -38.2 | -0.481 | -1.002 | 0.058 | 0.082 |
|  | ver | -40.8 | -0.524 | -0.970 | -0.072 | 0.182 |
| (Predators-ants):chewers ratio | ant | -36.3 | -0.451 | -1.041 | 0.143 | 0.134 |
|  | ver | -7.2 | -0.075 | -0.570 | 0.415 | 0.149 |
| Plant damage | ant | 147.2 | 0.905 | 0.521 | 1.295 | **<0.001** |
|  | ver | 46.8 | 0.384 | 0.004 | 0.762 | **<0.001** |

**Figure S2.** Effect of absolute latitude and treatment (exclusion of ants in red, exclusion of vertebrate insectivores in blue) on response variables. Latitude had no significant effect on any of the measured response variables (A, D-L). However, there was significant interaction between the latitude and the treatment (ant or vertebrate exclosure) in case of arthropod species richness (B) and herbivore arthropods (C). Y-axes show natural log ratios (LRR = ln(exclosure/control)) of individual response variables. Absolute latitude was used in models as well as plotted on x-axis. Elevation was used in all models as the first covariate. Effect size values above zero indicated that the absence of predators (treatment) was more harmful to plants than control (presence of predators), as the abundances of arthropods in exclosures increased and caused potentially higher herbivory damage to plants. Complete results of all models are in Table S2.

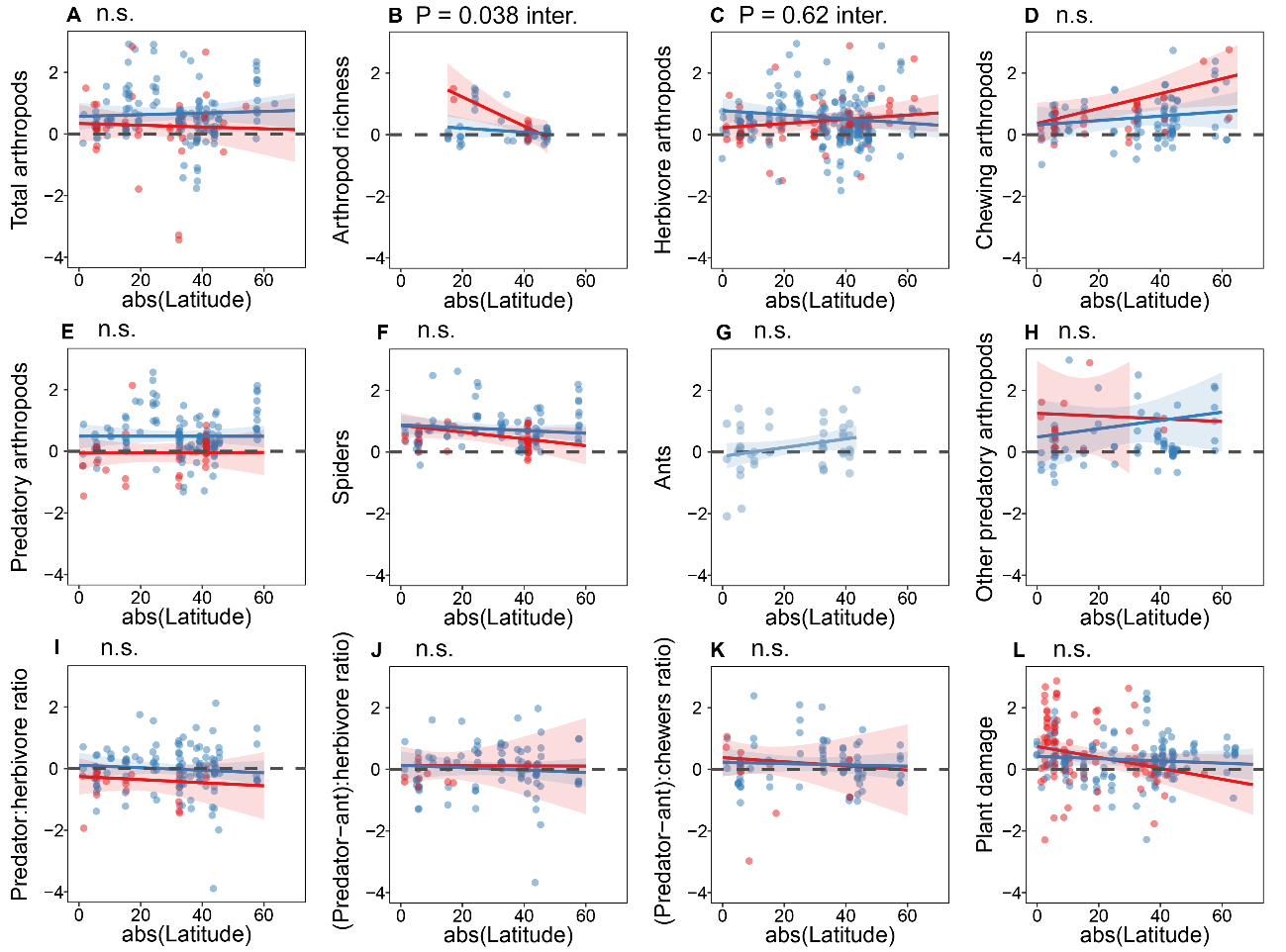

**Table S3**. Results of the models considering the effect of elevation (a) and latitude (b) and treatment on response variables. Latitude was used as covariate in models for elevation and elevation was used as covariate in models for latitude. Respective results are plotted in Fig. S2 and Figure 4.

| Models investigating effect of: | ELEVATION | | | | LATITUDE | | |
| --- | --- | --- | --- | --- | --- | --- | --- |
|  |  | Chisq | Df | P |  | Chisq | P |
| Total arthropods | Latitude | 0.059 | 1 | 0.807 | Elevation | 0.768 | 0.381 |
|  | *Elevation* | 0.565 | 1 | 0.452 | *Latitude* | 0.060 | 0.807 |
|  | Treatment | 4.290 | 1 | 0.038 | Treatment | 4.264 | 0.039 |
|  | *Elevation:Treatment* | 0.357 | 1 | 0.550 | *Latitude:Treatment* | 0.234 | 0.628 |
| Arthropod richness | Latitude | 0.505 | 1 | 0.477 | Elevation | 0.108 | 0.743 |
|  | *Elevation* | 0.003 | 1 | 0.958 | *Latitude* | 2.503 | 0.114 |
|  | Treatment | 2.910 | 1 | 0.088 | Treatment | 2.904 | 0.088 |
|  | *Elevation:Treatment* | 4.410 | 1 | **0.036** | *Latitude:Treatment* | 4.294 | **0.038** |
| Herbivore arthropods | Latitude | 0.361 | 1 | 0.548 | Elevation | 0.519 | 0.471 |
|  | *Elevation* | 0.429 | 1 | 0.513 | *Latitude* | 0.355 | 0.551 |
|  | Treatment | 1.310 | 1 | 0.252 | Treatment | 1.272 | 0.259 |
|  | *Elevation:Treatment* | 3.124 | 1 | **0.077** | *Latitude:Treatment* | 3.491 | **0.062** |
| Chewing arthropods | Latitude | 1.581 | 1 | 0.209 | Elevation | 1.802 | 0.179 |
|  | *Elevation* | 2.616 | 1 | 0.106 | *Latitude* | 2.831 | **0.093** |
|  | Treatment | 4.224 | 1 | 0.040 | Treatment | 3.970 | 0.046 |
|  | *Elevation:Treatment* | 5.905 | 1 | **0.015** | *Latitude:Treatment* | 1.970 | 0.16 |
| Predatory arthropods | Latitude | 0.008 | 1 | 0.930 | Elevation | 7.231 | 0.007 |
|  | Duration | 7.166 | 1 | 0.007 | Duration | 6.742 | 0.009 |
|  | *Elevation* | 7.723 | 1 | **0.005** | *Latitude* | 0.000 | 0.997 |
|  | Treatment | 16.237 | 1 | <0.001 | Treatment | 16.045 | <0.001 |
|  | *Elevation:Treatment* | 1.131 | 1 | 0.288 | *Latitude:Treatment* | 0.003 | 0.957 |
| Spiders | Latitude | 1.235 | 1 | 0.266 | Elevation | 12.443 | <0.001 |
|  | *Elevation* | 10.959 | 1 | **<0.001** | *Latitude* | 1.592 | 0.207 |
|  | Treatment | 1.669 | 1 | 0.196 | Treatment | 1.361 | 0.243 |
|  | *Elevation:Treatment* | 1.857 | 1 | 0.173 | *Latitude:Treatment* | 0.842 | 0.359 |
| Ants (only for vert. exclosures) | Latitude | 4.609 | 1 |  |  |  |  |
|  | *Elevation* | 2.132 | 1 |  |  |  |  |
| Other predatory arthropods | Latitude | 0.386 | 1 | 0.535 | Elevation | 0.118 | 0.731 |
|  | *Elevation* | 0.272 | 1 | 0.602 | *Latitude* | 0.430 | 0.512 |
|  | Treatment | 0.759 | 1 | 0.384 | Treatment | 0.798 | 0.372 |
|  | *Elevation:Treatment* | 0.137 | 1 | 0.711 | *Latitude:Treatment* | 0.174 | 0.677 |
| Predatore:herbivore ratio | Latitude | 0.617 | 1 | 0.432 | Elevation | 1.840 | 0.175 |
|  | Duration | 8.679 | 1 | 0.003 | Duration | 8.529 | 0.003 |
|  | *Elevation* | 2.396 | 1 | 0.122 | *Latitude* | 0.515 | 0.473 |
|  | Treatment | 4.624 | 1 | 0.032 | Treatment | 4.528 | 0.033 |
|  | *Elevation:Treatment* | 0.008 | 1 | 0.928 | *Latitude:Treatment* | 0.005 | 0.942 |
| (Predators-ants):herbivore ratio | Latitude | 0.400 | 1 | 0.527 | Elevation | 4.069 | 0.044 |
|  | Duration | 7.631 | 1 | 0.006 | Duration | 8.503 | 0.004 |
|  | *Elevation* | 4.189 | 1 | **0.041** | *Latitude* | 0.287 | 0.592 |
|  | Treatment | 0.027 | 1 | 0.869 | Treatment | 0.037 | 0.847 |
|  | *Elevation:Treatment* | 0.264 | 1 | 0.608 | *Latitude:Treatment* | 0.046 | 0.830 |
| (Predators-ants):chewers ratio | Latitude | 0.174 | 1 | 0.676 | Elevation | 2.371 | 0.124 |
|  | Duration | 3.462 | 1 | 0.063 | Duration | 5.957 | 0.015 |
|  | *Elevation* | 1.793 | 1 | 0.181 | *Latitude* | 0.149 | 0.699 |
|  | Treatment | 0.252 | 1 | 0.616 | Treatment | 0.232 | 0.630 |
|  | Duration:Treatment | 8.234 | 1 | 0.004 | Duration:Treatment | 15.432 | <0.001 |
|  | *Elevation:Treatment* | 0.573 | 1 | 0.449 | *Latitude:Treatment* | 0.093 | 0.761 |
| Plant damage | Latitude | 0.931 | 1 | 0.335 | Elevation | 0.696 | 0.404 |
|  | Duration | 3.115 | 1 | 0.077 | Duration | 4.149 | 0.042 |
|  | *Elevation* | 0.880 | 1 | 0.348 | *Latitude* | 2.133 | 0.144 |
|  | Treatment | 0.015 | 1 | 0.903 | Treatment | 0.175 | 0.676 |
|  | Duration:Treatment | 3.662 | 1 | 0.056 | Duration:Treatment | 4.770 | 0.029 |
|  | *Elevation:Treatment* | 4.687 | 1 | **0.030** | *Latitude:Treatment* | 1.943 | 0.163 |

**Figure S3.** Effect of mean NDVI of productive season (a) and median NDVI of all monthly values (b) and treatment on response variables. Y-axes show natural log ratios (LRR = ln(exclosure/control)) of individual response variables. Complete results of all models are in Table S4.

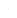

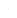

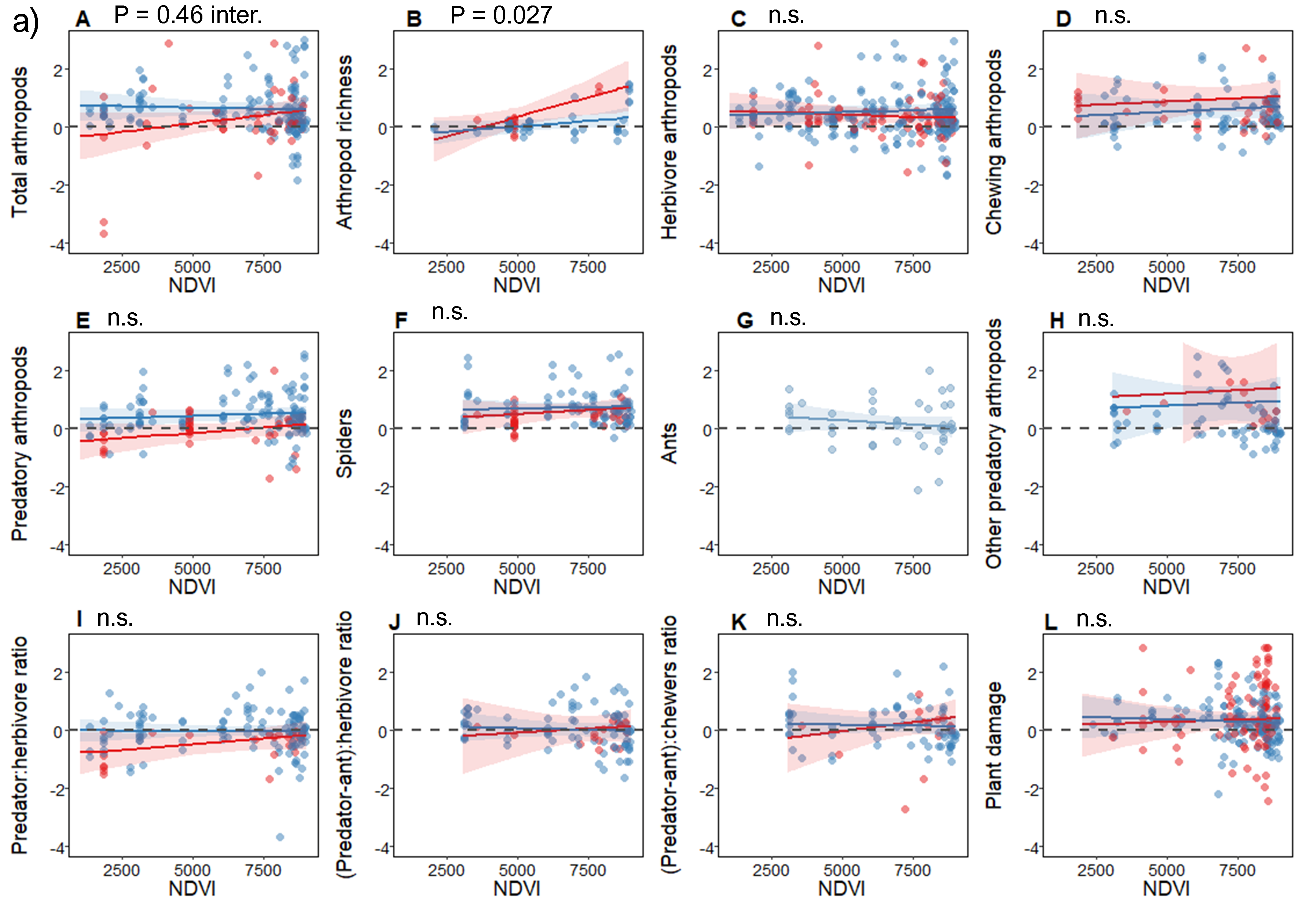

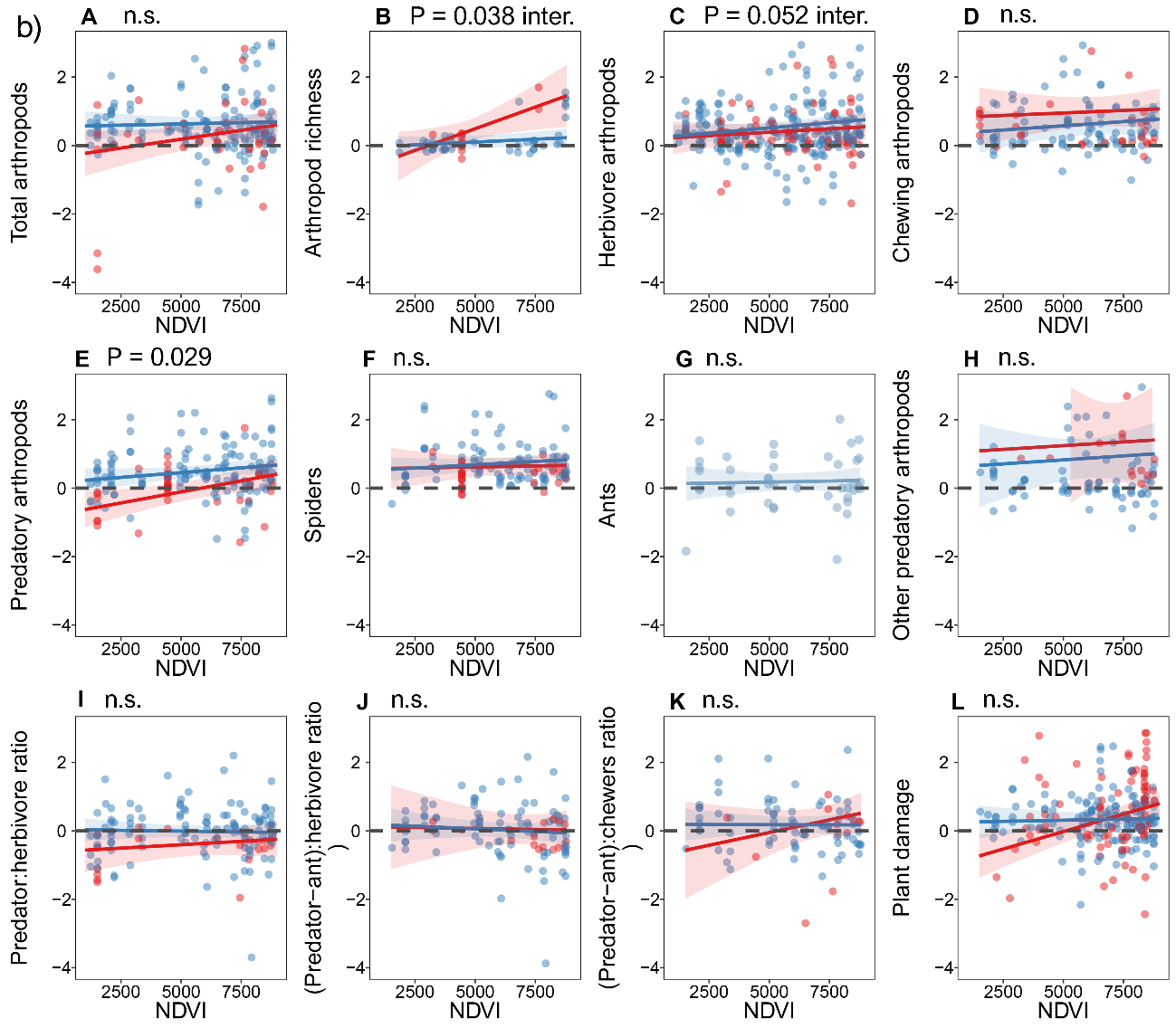

**Table S4**. Results of the models considering the effect of precipitation (a), mean NDVI of productive season (b), median NDVI of all monthly values (c) and treatment on response variables. Lines with focal factors are shaded in grey. Significant, and nearly significant, values for focal factors are marked in bold. Duration is used as covariate in models for responses which were earlier found to be affected by the duration of the experiment (refer to Fig. S1 and Table S1). Respective results are plotted in Fig. 5 and Fig. S3.

| Models investigating effect of: | a) PRECIPITATION | | | b) Mean NDVI of productive season | | | c) Median NDVI of all months | | |
| --- | --- | --- | --- | --- | --- | --- | --- | --- | --- |
|  |  | Chisq | P |  | Chisq | P |  | Chisq | P |
| Total arthropods | Precip | 0.060 | 0.807 | NDVI | 0.029 | 0.865 | NDVI | 1.015 | 0.314 |
|  | Treatment | 4.549 | 0.033 | Treatment | 4.514 | 0.034 | Treatment | 4.633 | 0.032 |
|  | Precip:Treatment | 3.232 | **0.072.** | NDVI:Treatment | 3.969 | **0.046** | NDVI:Treatment | 2.153 | 0.142 |
| Arthropod richness | Precip | 2.418 | 0.120 | NDVI | 4.893 | **0.027** | NDVI | 2.546 | 0.111 |
|  | Treatment | 2.982 | 0.084 | Treatment | 4.041 | 0.044 | Treatment | 3.560 | 0.059 |
|  | Precip:Treatment | 4.831 | **0.028** | NDVI:Treatment | 2.872 | **0.090.** | NDVI:Treatment | 4.218 | **0.040** |
| Herbivore arthropods | Precip | 0.198 | 0.657 | NDVI | 0.207 | 0.649 | NDVI | 3.784 | **0.052.** |
|  | Treatment | 1.260 | 0.262 | Treatment | 1.193 | 0.275 | Treatment | 1.214 | 0.271 |
|  | Precip:Treatment | 4.858 | **0.028** | NDVI:Treatment | 1.294 | 0.255 | NDVI:Treatment | 0.179 | 0.672 |
| Chewing arthropods | Precip | 0.053 | 0.819 | NDVI | 0.540 | 0.462 | NDVI | 0.645 | 0.422 |
|  | Treatment | 2.242 | 0.134 | Treatment | 2.263 | 0.133 | Treatment | 2.210 | 0.137 |
|  | Precip:Treatment | 1.463 | 0.227 | NDVI:Treatment | 0.000 | 0.991 | NDVI:Treatment | 0.044 | 0.834 |
| Predatory arthropods | Duration | 6.991 | 0.008 | Duration | 6.907 | 0.008 | Duration | 7.318 | 0.007 |
|  | Precip | 2.105 | 0.147 | NDVI | 0.979 | 0.322 | NDVI | 4.731 | **0.029** |
|  | Treatment | 16.008 | <0.001 | Treatment | 15.341 | <0.001 | Treatment | 15.484 | <0.001 |
|  | Precip:Treatment | 0.320 | 0.572 | NDVI:Treatment | 0.852 | 0.356 | NDVI:Treatment | 2.490 | 0.115 |
| Spiders | Precip | 4.836 | **0.028** | NDVI | 0.398 | 0.528 | NDVI | 1.176 | 0.278 |
|  | Treatment | 0.801 | 0.371 | Treatment | 0.790 | 0.374 | Treatment | 0.769 | 0.380 |
|  | Precip:Treatment | 0.000 | 0.998 | NDVI:Treatment | 0.272 | 0.602 | NDVI:Treatment | 0.195 | 0.659 |
| Ants (only for vert. exclosures) | Precip | 1.037 | 0.309 | NDVI | 1.086 | 0.297 | NDVI | 0.306 | 0.579 |
| Other predatory arthropods | Precip | 0.600 | 0.438 | NDVI | 0.086 | 0.769 | NDVI | 0.151 | 0.698 |
|  | Treatment | 0.656 | 0.418 | Treatment | 0.553 | 0.457 | Treatment | 0.481 | 0.488 |
|  | Precip:Treatment | 0.032 | 0.858 | NDVI:Treatment | 0.001 | 0.977 | NDVI:Treatment | 0.000 | 0.993 |
| Predatore:herbivore ratio | Duration | 8.482 | 0.004 | Duration | 5.835 | 0.015 | Duration | 8.060 | 0.005 |
|  | Precip | 1.114 | 0.291 | NDVI | 0.070 | 0.792 | NDVI | 0.004 | 0.950 |
|  | Treatment | 4.595 | 0.032 | Treatment | 0.059 | 0.807 | Treatment | 4.302 | 0.038 |
|  | Precip:Treatment | 1.118 | 0.290 | NDVI:Treatment | 0.309 | 0.578 | NDVI:Treatment | 0.673 | 0.412 |
| (Predators-ants):herbivore ratio | Duration | 7.577 | 0.006 | Duration | 5.835 | 0.018 | Duration | 6.212 | 0.013 |
|  | Precip | 1.027 | 0.311 | NDVI | 0.070 | 0.792 | NDVI | 0.402 | 0.526 |
|  | Treatment | 0.015 | 0.902 | Treatment | 0.059 | 0.807 | Treatment | 0.054 | 0.816 |
|  | Precip:Treatment | 0.198 | 0.656 | NDVI:Treatment | 0.309 | 0.578 | NDVI:Treatment | 0.031 | 0.861 |
| (Predators-ants):chewers ratio | Duration | 5.937 | 0.015 | Duration | 4.981 | 0.026 | Duration | 5.374 | 0.021 |
|  | Precip | 0.748 | 0.387 | NDVI | 0.004 | 0.951 | NDVI | 0.053 | 0.819 |
|  | Treatment | 0.222 | 0.638 | Treatment | 0.102 | 0.749 | Treatment | 0.142 | 0.706 |
|  | Duration:Treatment | 14.796 | <0.001 | Duration:Treatment | 15.203 | <0.001 | Duration:Treatment | 15.325 | <0.001 |
|  | Precip:Treatment | 2.195 | 0.138 | NDVI:Treatment | 1.012 | 0.314 | NDVI:Treatment | 1.479 | 0.224 |
| Plant damage | Duration | 4.615 | 0.032 | Duration | 4.711 | 0.029 | Duration | 4.849 | 0.028 |
|  | Precip | 0.805 | 0.370 | NDVI | 0.040 | 0.842 | NDVI | 6.234 | **0.013** |
|  | Treatment | 0.065 | 0.799 | Treatment | 0.166 | 0.684 | Treatment | 0.061 | 0.805 |
|  | Duration:Treatment | 4.065 | 0.044 | Duration:Treatment | 3.832 | 0.050 | Duration:Treatment | 4.143 | 0.042 |
|  | Precip:Treatment | 0.080 | 0.778 | NDVI:Treatment | 0.278 | 0.598 | NDVI:Treatment | 8.068 | **0.005** |

**Figure S4**. Effect of habitat (agricultural vs. natural) and treatment (exclusion of ants in red, exclusion of vertebrate insectivores in blue) on response variables. Type of the habitat did not have significant effect on most of the surveyed response variables, except on abundances of herbivore arthropods (C) and spiders (F). Abundances of herbivore arthropods increased significantly more in agricultural habitats after exclosure of vertebrate insectivores, but they did not increase when ants were excluded (C). Similar trend was observed in natural habitats (C). While abundances of spiders increase the most in agricultural habitats after the vertebrate insectivores were removed, there was no detectable difference in their increase in agricultural and natural habitats after ants were removed (F). Y-axes show natural log ratios (LRR = ln(exclosure/control)) of individual response variables. Complete results of all models are in Table S5.

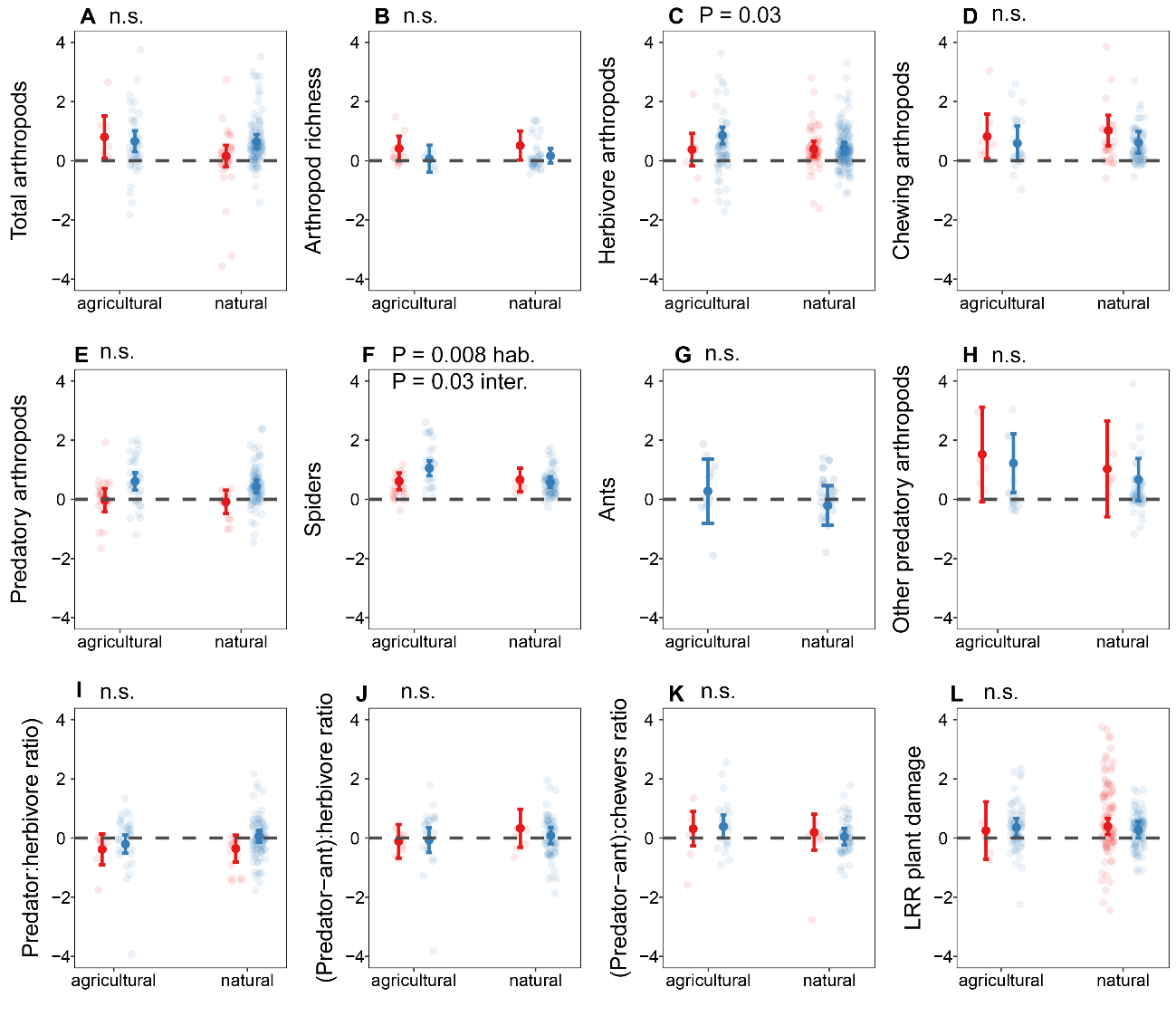

**Figure S5**. Effect of growth from (mature tree vs. sapling) and treatment (exclusion of ants in red, exclusion of vertebrate insectivores in blue) on response variables. Arthropod richness (B) and abundances of other predatory arthropods (H) increased significantly more on saplings than on mature trees in both treatments. On the other hand, abundances of ants increased significantly more on mature trees than on saplings in vertebrate exclosures (G). The abundances of chewing arthropods (D) and predatory arthropods (E) and the ratio between predatory arthropods without ants and chewers interacted with growth form. Abundances of herbivore arthropods increased significantly more in agricultural habitats after exclosure of vertebrate Y-axes show natural log ratios (LRR = ln(exclosure/control)) of individual response variables. Complete results of all models are in Table S5.

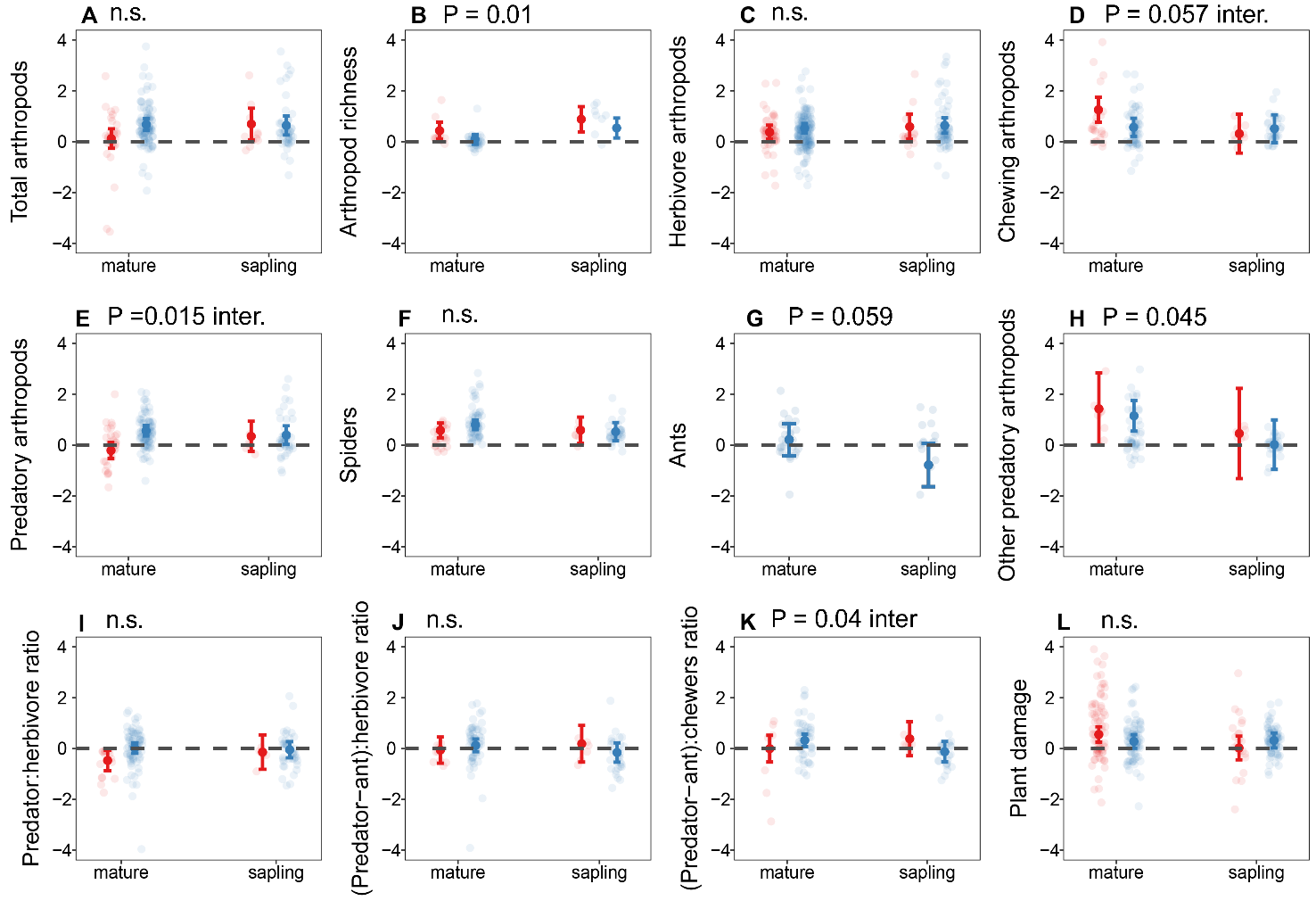

**Table S5**. Results of the models considering the effect of habitat (a) and plant form (b) and treatment on response variables. Respective results are visually presented in Fig. S4 and Fig. S5.

| Models investigating effect of: | HABITAT | | | PLANT GROWTH FORM | | |
| --- | --- | --- | --- | --- | --- | --- |
|  |  | Chisq | P |  | Chisq | P |
| Total arthropods | Habitat | 0.415 | 0.520 | Form | 0.139 | 0.710 |
|  | Treatment | 4.293 | 0.038 | Treatment | 4.442 | 0.035 |
|  | Habitat:Treatment | 2.152 | 0.142 | Form:Treatment | 2.634 | 0.105 |
| Arthropod richness | Habitat | 0.147 | 0.701 | Form | 5.413 | **0.010** |
|  | Treatment | 2.688 | 0.101 | Treatment | 3.915 | 0.040 |
|  | NA | | |  |  |  |
| Herbivore arthropods | Habitat | 4.563 | **0.033** | Form | 0.568 | 0.451 |
|  | Treatment | 0.829 | 0.363 | Treatment | 1.120 | 0.290 |
|  | Habitat:Treatment | 1.962 | 0.161 | Form:Treatment | 0.139 | 0.709 |
| Chewing arthropods | Habitat | 0.074 | 0.786 | Form | 0.911 | 0.340 |
|  | Treatment | 2.242 | 0.134 | Treatment | 2.342 | 0.126 |
|  | Habitat:Treatment | 0.114 | 0.735 | Form:Treatment | 3.606 | **0.057** |
| Predatory arthropods | Duration | 6.029 | 0.014 | Duration | 6.626 | 0.010 |
|  | Habitat | 0.703 | 0.402 | Form | 0.242 | 0.623 |
|  | Treatment | 16.520 | <0.001 | Treatment | 17.122 | <0.001 |
|  | Habitat:Treatment | 0.159 | 0.690 | Form:Treatment | 6.031 | **0.014** |
| Spiders | Habitat | 7.013 | **0.008** | Form | 1.801 | 0.180 |
|  | Treatment | 4.292 | 0.038 | Treatment | 1.280 | 0.258 |
|  | Habitat:Treatment | 4.713 | **0.030** | Form:Treatment | 1.412 | 0.235 |
| Ants (only for vert. exclosures) | Habitat | 0.584 | 0.445 | Form | 3.573 | 0.059 |
| Other predatory arthropods | Habitat | 0.894 | 0.344 | Form | 4.013 | **0.045** |
|  | Treatment | 0.316 | 0.574 | Treatment | 0.369 | 0.544 |
|  | Habitat:Treatment | 0.003 | 0.959 | Form:Treatment | 0.020 | 0.887 |
| Predator:herbivore ratio | Duration | 8.892 | 0.003 | Duration | 7.733 | 0.005 |
|  | Habitat | 1.702 | 0.192 | Form | 0.019 | 0.890 |
|  | Treatment | 3.387 | 0.066 | Treatment | 4.317 | 0.037 |
|  | Habitat:Treatment | 0.427 | 0.513 | Form:Treatment | 1.017 | 0.313 |
| (Predators-ants):herbivore ratio | Duration | 6.719 | 0.010 | Duration | 6.648 | 0.009 |
|  | Habitat | 0.727 | 0.394 | Form | 0.897 | 0.344 |
|  | Treatment | 0.203 | 0.652 | Treatment | 0.014 | 0.906 |
|  | Habitat:Treatment | 0.418 | 0.517 | Form:Treatment | 1.422 | 0.233 |
| (Predators-ants):chewers ratio | Duration | 5.664 | 0.017 | Duration | 6.282 | 0.012 |
|  | Habitat | 1.747 | 0.186 | Form | 2.019 | 0.155 |
|  | Treatment | 0.369 | 0.544 | Treatment | 0.408 | 0.523 |
|  | Duration:Treatment | 14.998 | <0.001 | Duration:Treatment | 15.939 | <0.001 |
|  | Habitat:Treatment | 0.269 | 0.604 | Form:Treatment | 4.208 | **0.040** |
| Plant damage | Duration | 4.450 | 0.035 | Duration | 3.577 | 0.058 |
|  | Habitat | 0.058 | 0.810 | Form | 0.562 | 0.454 |
|  | Treatment | 0.073 | 0.787 | Treatment | 0.069 | 0.793 |
|  | Duration:Treatment | 3.788 | 0.052 | Duration:Treatment | 2.798 | 0.094 |
|  | Habitat:Treatment | 0.156 | 0.693 | Form:Treatment | 3.106 | **0.077** |
